# Community responses outweigh species-level plasticity in shaping future marine forests

**DOI:** 10.64898/2026.11.16.766521

**Authors:** Ingrid Graf, Anaïs Barrera, Emma Cebrian, Cristina Galobart, Giovanni Vecchiatti, Marcos Fontela, Núria Teixidó, Luisa Mangialajo, Jana Verdura, Jéremy Carlot, Steeve Comeau, Jean-Pierre Gattuso, Samir Alliouane, Valentina Esposito, Jordi Boada

**Author notes:** Corresponding author: Jordi Boada.

## Abstract

Anthropogenic ocean warming (OW) and acidification (OA) are progressively reshaping marine ecosystems, yet ecosystem-level evidence remains limited. Marine forests are foundational ecosystems that support highly productive and diverse communities. Understanding how macroalgal communities will assemble, perform and persist in future ocean conditions requires integrated evidence capturing concurrent shifts in primary productivity, species interactions, and associated macroalgal community responses. Here, we provide a multi-level approach combining laboratory experiments with field surveys to assess how Mediterranean marine macroalgal forests will respond to future warming and acidification. We used local populations from natural CO₂ vents in Panarea (Aeolian Islands, Italy) and present-day pH conditions of the emblematic forest-forming species *Ericaria balearica* to quantify multi-level responses to future climate drivers. Specifically, we crossed temperature (present day, T+2, T+4 °C), pH (present day pH_T_ ∼ 7.9, low pH_T_ ∼7.6), and origin population (ambient vs CO₂ vent) in laboratory experiments, where we measured changes in *Ericaria* biomass and metabolism, as well as herbivory by the sea urchin *Paracentrotus lividus*. We also assessed *in situ* community structure by quantifying species-specific percent cover and biomass along a pH gradient. In natural CO₂ vents, *E. balearica* biomass was greater than at ambient sites, and associated macroalgal community composition differed between pH zones. In the laboratory, across the simulated spring–summer warming period, *E. balerarica* biomass declined similarly in all treatments with no important effect of pH; a marginal signal suggested that vent-origin individuals retained more biomass only under present-day temperature. Sea urchin consumption was reduced under low pH and showed a weaker response to temperature. Overall, we found no clear direct impact of OW×OA on *E. balearica* performance but significant influence of OA on herbivory and the associated macroalgal community, indicating that ecological processes outweigh the effects at the individual or population level.

## Introduction

Understanding how key ecosystems will respond to rapid, climate-driven changes in ocean conditions remains a major challenge. Rising greenhouse gas concentrations in the atmosphere are driving global ocean warming (OW). Concurrently, the ocean absorbs about 25% of anthropogenic CO₂, shifting the marine carbonate system and triggering widespread ocean acidification (OA). Under high-emissions scenarios (SSP5-8.5), sea-surface temperatures are projected to increase by roughly 3.47°C relative to pre-industrial baselines by 2100, while continued CO₂ uptake will further lower pH by –0.44 and carbonate saturation states (Gattuso et al., 2015; Kwiatkowski et al., 2020). These rapidly changing conditions are already altering marine ecosystems globally, driving changes in species distributions and persistent functional shifts, such as transitions from kelp forests to urchin barrens and from structurally complex to simplified coral reef assemblages (Wernberg et al., 2016; Hughes et al., 2018; Rogers-Bennett & Catton, 2019; Armstrong McKay et al., 2022). Consequently, resolving the ecological mechanisms that drive species performance and overall resilience is critical to predict ecosystem trajectories under future climate scenarios. However, most of the scientific evidence is based on single species-single driver or on the modeling of extreme events (e.g., marine heatwaves) rather than understanding species sensitivity and community responses to expected changing environmental conditions.

In the ocean, foundation seagrasses and canopy-forming macroalgae play a disproportionate role in supporting biodiversity and primary production, protecting shorelines from erosion, improving water quality, and in some contexts buffering climate change through carbon sequestration and local pH amelioration (Olafsson, 2016; Ricart et al., 2021; Pessarrodona et al., 2022). These habitats have undergone widespread canopy declines and regime shifts driven by global warming and local stressors such as eutrophication, turbidity, habitat modification, and overgrazing (Krumhansl et al., 2016; Wernberg et al., 2016; Boada et al., 2017;), yet there is still limited capacity to evaluate their ecological status under future environmental regimes.

Mounting evidence highlights the catastrophic consequences of OW on kelp and seagrasses. Particularly, marine heatwaves (MHWs) have triggered large-scale range contractions across both temperate and tropical regions, shifting vibrant marine forests into highly stable, simplified ecosystem states (Wernberg et al., 2016; Arias-Ortiz et al., 2018; Straub et al., 2019; Verdura et al., 2021). For canopy-forming *Fucales* algae like *Cystoseira s.l.*, thermal stress commonly impairs survival, growth, and reproductive performance while their photophysiology severely deteriorates as temperatures approach upper thermal tolerance limits (Hernández et al., 2018; Savva et al., 2018; Capdevila et al., 2019; Verdura et al., 2021).

In contrast, elevated *pCO₂*associated with OA can positively affect many non-calcifying foundational macroalgae and seagrasses by enhancing photosynthetic capacity and growth (Koch et al., 2013; Kroeker et al., 2013; Celis-Plá et al., 2015). By comparison, OA can impair biomineralization and promote net dissolution in calcareous and crustose coralline algae (Johnson et al., 2014; Cornwall et al., 2022). These responses are also modulated by seasonality, with their magnitude and direction depending on the physiological state of the organism and prevailing environmental conditions (Baggini et al., 2014; Graiff et al., 2015; Truong et al., 2024). Thus, OA effects extend beyond individual growth and persistence to influence reproductive timing and success, with potential consequences for population and community dynamics.

Yet, the long-term resilience of these ecosystems depends on more than just the standalone physiological effects of OW and OA. They are also fundamentally shaped by ecological interactions with other organisms, including herbivory, competition, and facilitation. OW, for instance, can elevate feeding rates, the movement, and metabolic demands of consumers, potentially intensifying herbivory and altering canopy–consumer balances (Pagès et al., 2018). Concurrently, OA can reshape community structures by negatively impacting calcifying algae and herbivores while favoring non-calcifying competitors, fundamentally altering the strength of these ecological relationships (Kroeker et al., 2013). Despite these complex dynamics, studies quantifying the interactive effects of OW and OA on macroalgal communities remain sparse, limiting our understanding of future ecosystem trajectories (but see Koch et al., 2013; Celis-Plá et al., 2017; Monserrat et al., 2022).

Realistically projecting the future functioning of marine forests requires a multi-level assessment of responses to OW and OA together, in form of future ocean environmental conditions simulations complementary to responses of organisms to temperature and pH extremes. This type of experimentation has not been prioritized probably due to the complexity of their implementation and the expectation that the responses observed are limited compared to the response to extreme stress. Here, we address this gap by investigating future ecosystem functioning using as a case study, the species *Ericaria balearica*, a forest-forming macroalga of the order Fucales. To this end, we implemented an integrative approach that combined 1) physiological and metabolic rate assessments, 2) plant-herbivore interactions, and 3) community assessments.

We took advantage of populations *of E. balearica* naturally occurring in natural CO_2_ vent systems, in Panarea, Italy, which serve as natural analogs of projected OA. These systems allow us to evaluate the long-term ecological effects of chronic exposure to low-pH conditions (Porzio et al., 2017, Teixidó et al 2024). In dedicated expeditions, we quantified the natural populations associated to these CO_2_ systems and compared them with those occurring under ambient pH conditions. Furthermore, we used individuals of these local populations in a series of manipulative laboratory experiments mimicking OW and OA regimes to project their future functioning. We hypothesized that 1) OA enhances *E. balearica* performance via carbon fertilization and that the population historically exposed to low-pH further benefits from increased CO_2_ availability, 2) a decrease in herbivory pressure due to reduced performance of calcifying herbivores, and 3) shifts in the associated macroalgal community.

## Methods

### Study site and sampling

The Panarea Island (Aeolian archipelago, Italy) hosts one of the largest hydrothermal seep complexes in the Mediterranean Sea, including the crater off Bottaro Islet where a well-defined pH gradient (pH_T_ ∼ 7.40-7.99) extends from the CO_2_ caldera to the surrounding seabed (Esposito et al., 2022). Around the crater rim and adjacent bottoms, dense stands of *E. balearica* structure the shallow infralittoral habitat. *E. balearica* is an epilithic brown alga inhabiting the infralittoral zone (0-25m depth), characterized by a perennial, caespitose thallus. Its growth displays marked seasonal dynamics, with its peak during spring and summer. As a key canopy-forming foundation species, it provides critical three-dimensional habitat, high primary productivity, and nursery grounds for coastal Mediterranean biodiversity (Hoffman et al., 1992; Mannino et al., 2026).

To evaluate the physiological responses of *E. balearica* to projected OW and OA conditions, we conducted a controlled laboratory experiment at the Centre d’Estudis Avançats de Blanes (CEAB-CSIC) using populations exposed to contrasting pH regimes near Panarea. Individuals were collected near Bottaro Islet (CO₂ Vent site; 38°38′13″N, 15°06′36″E,) and near Dattilo (ambient pH site; 38°38′20.37″N, 15°05′55.12″E,), which has similar hydrodynamic and substrate conditions but not venting activity. Samples were collected at 7 m depth at both sites by SCUBA diving the 3rd of May 2025 (approximately 60 individuals per treatment), when sea-surface temperature was 19.1 °C (Copernicus Marine Service, 2025; product SST_MED_SST_L4_NRT_OBSERVATIONS_010_004). Individuals were cleaned to remove epiphytes and organic matter, packed in plastic bags and transported in cold and dark conditions to the CEAB-CSIC laboratory in 48h.

### Manipulative laboratory experiments

#### (i) Seaweed performance

To test seaweed performance under future conditions, we used a fully factorial design crossing three temperatures and two pH levels based on IPCC end of century projections (Kwiatkowski et al., 2020; Fox-Kemper et al., 2021). Temperature treatments tracked the local sea surface temperature (SST) cycle in Panarea, increasing progressively throughout the experiment to mimic natural seasonal warming. The three temperature treatments consisted of present-day SST, T+2 °C to represent moderate warming (SSP2-4.5), and +4 °C to represent high warming (SSP5-8.5). After the acclimation period, the water temperature was gradually increased by 0.2°C daily to simulate the natural seasonal progression from spring to summer. pH_T_ was held either at ambient pH value of ∼7.9 (present day) or reduced to ∼7.60 to simulate projected acidification by 2100 under the scenario SSP5-8.5. We crossed these factors with origin (ambient site vs. CO₂-vent site), yielding 12 treatment combinations (2 local populations × 2 pH conditions × 3 ramping temperatures). Each treatment had eight replicates, totaling 96 *E. balearica* individuals. Samples were randomly assigned to one of the treatments and acclimated for 15 days. The temperature during the acclimation period gradually increased by 0.5 C per day until treatment temperatures were reached to prevent thermic shock. Six tanks (160 L) were used as temperature-controlled water baths, 2 tanks per temperature treatment. Each tank contained eight aquaria (12.6 L), resulting in a total of 48 aquaria. Within each tank, half of the aquaria were assigned to a pH treatment. One individual from each local population was randomly assigned to every aquarium. Individuals from the CO_2_ vent were tagged for identification. The 47-day experiment was conducted under natural ambient light conditions.

### Carbonate chemistry

Low-pH seawater was prepared in header tanks by mixing CO₂ gas into ambient seawater using a pressurized CO₂ system until the target pH was reached (pH_T_ 7.6). The system operated in an open circuit, with seawater continuously delivered to each aquarium with individual tubes. Water flow was set to allow complete water renewal on a daily basis within aquaria. Temperature was continuously monitored using GHL Profilux 4 digital temperature sensor (PL-1115, resolution 0.1°C) and HOBO Pendant MX Temperature/Light Data Loggers (MX2202, temperature resolution 0.04 °C). pH was monitored using GHL Profilux 4 pH electrode (PL-0071, resolution 0.01 pH) calibrated with GHL pH 7.0 (PL-0075) and 9.2 (PL-2343) buffer solutions. Water samples were taken at three points during the experimental period to measure total alkalinity (A_T_). For each treatment, four 125 mL unfiltered water samples were collected in plastic bottles and spiked with 62 µL of saturated HgCl₂ solution to halt biological activity. Bottles were sealed and stored in the dark at room temperature until their analysis eight months later (March 2026). A_T_ was analyzed by single point titration (Pérez & Fraga, 1987b) and compared to certified reference materials (CRM #218 ±3.6 μmol kg^-1^, Dickson et al., 2007) following Fontela et al., (2023). The analysis was adjusted to the available sample volume with a gravimetrically calibrated Knudsen pipette (50 mL) and the results have a measurement precision of ± 5 μmol kg−1. Measured pH values (NBS scale) were converted to the total scale (pH_T_) using the pHnbs2sws() and pHconv() functions of the R package *seacarb* v3.3.4 (Gattuso et al., 2026), which convert pH from the NBS to the seawater scale and subsequently to the total scale, accounting for temperature and salinity (38). A_T_ and pH_T_ were used to determine the remaining carbonate system variables (Table 1). Carbonate system parameters were calculated using K₁ and K₂ from Lueker et al. (2000), Kf from Pérez and Fraga (1987a), Ks from Dickson (1990), and total boron from Uppström (1974).

**Table 1.** Seawater physicochemical parameters across experimental treatments. Parameters include pH_T_, temperature (T), total alkalinity (A_T_), dissolved inorganic carbon (C_T_), and *p*CO₂. Mean pH_T_ was calculated by averaging hydrogen-ion concentrations and converting back to pH units, with variability expressed as the 25th–75th percentiles. Other parameters are presented as means ± SD (25th–75th percentiles).

| **pH condition** | **Temperature condition** | **Date** | **pH<sub>T</sub>** | **T (°C)** | **A<sub>T</sub> (µmol kg<sup>-1</sup>)** | **C<sub>T</sub> (µmol kg<sup>-1</sup>)** | ***p*CO<sub>2</sub> (µatm)** |
|---|---|---|---|---|---|---|---|
| Present-day | T | 03/06/2025 | 7.90 (7.89–7.90) | 21.51 ± 0.6 (20.93–22.09) | 2599 ± 29 (2579–2622) | 2370 ± 28 (2352–2392) | 670 ± 8 (665–676) |
| Present-day | T+2 | 03/06/2025 | 7.89 (7.88–7.89) | 23.84 ± 0.23 (23.63–24.06) | 2580 ± 23 (2568–2583) | 2338 ± 21 (2326–2340) | 675 ± 6 (671–675) |
| Present-day | T+4 | 03/06/2025 | 7.88 (7.88–7.89) | 26.95 ± 0.31 (26.72–27.03) | 2585 ± 24 (2571–2589) | 2320 ± 22 (2307–2323) | 688 ± 7 (684–689) |
| Present-day | T | 17/06/2025 | 7.85 (7.85–7.85) | 26 ± 0.22 (25.82–26.22) | 2614 ± 29 (2596–2622) | 2374 ± 27 (2357–2382) | 764 ± 9 (758–766) |
| Present-day | T+2 | 17/06/2025 | 7.85 (7.84–7.85) | 27.39 ± 0.21 (27.2–27.54) | 2593 ± 27 (2574–2602) | 2345 ± 26 (2328–2354) | 764 ± 8 (758–767) |
| Present-day | T+4 | 17/06/2025 | 7.84 (7.84–7.84) | 29.13 ± 0.22 (28.95–29.35) | 2609 ± 27 (2596–2618) | 2348 ± 25 (2335–2356) | 776 ± 8 (772–779) |
| Present-day | T | 07/07/2025 | 7.82 (7.79–7.83) | 28.71 ± 0.38 (28.27–29.01) | 2606 ± 27 (2591–2615) | 2363 ± 25 (2348–2371) | 828 ± 9 (823–831) |
| Present-day | T+2 | 07/07/2025 | 7.81 (7.79–7.83) | 30.26 ± 0.77 (30.18–30.8) | 2605 ± 25 (2591–2614) | 2351 ± 23 (2338–2359) | 835 ± 8 (830–838) |
| Present-day | T+4 | 07/07/2025 | 7.81 (7.78–7.82) | 32.23 ± 0.35 (31.95–32.47) | 2718 ± 43 (2703–2733) | 2441 ± 40 (2427–2455) | 881 ± 14 (876–886) |
| low pH | T | 03/06/2025 | 7.54 (7.52–7.58) | 21.51 ± 0.6 (20.93–22.09) | 2573 ± 8 (2570–2576) | 2503 ± 8 (2500–2505) | 1638 ± 5 (1637–1640) |
| low pH | T+2 | 03/06/2025 | 7.55 (7.52–7.59) | 23.84 ± 0.23 (23.63–24.06) | 2669 ± 32 (2651–2680) | 2584 ± 32 (2566–2594) | 1700 ± 21 (1688–1706) |
| low pH | T+4 | 03/06/2025 | 7.54 (7.50–7.59) | 26.95 ± 0.31 (26.72–27.03) | 2580 ± 22 (2567–2586) | 2481 ± 22 (2469–2488) | 1673 ± 15 (1665–1678) |
| low pH | T | 17/06/2025 | 7.56 (7.54–7.57) | 26 ± 0.22 (25.82–26.22) | 2625 ± 51 (2589–2652) | 2523 ± 50 (2488–2549) | 1624 ± 32 (1601–1641) |
| low pH | T+2 | 17/06/2025 | 7.56 (7.54–7.59) | 27.39 ± 0.21 (27.2–27.54) | 2594 ± 20 (2583–2603) | 2484 ± 19 (2474–2493) | 1602 ± 13 (1595–1608) |
| low pH | T+4 | 17/06/2025 | 7.55 (7.54–7.57) | 29.13 ± 0.22 (28.95–29.35) | 2611 ± 47 (2584–2628) | 2494 ± 45 (2468–2510) | 1649 ± 30 (1632–1660) |
| low pH | T | 07/07/2025 | 7.55 (7.54–7.58) | 28.71 ± 0.38 (28.27–29.01) | 2646 ± 25 (2633–2657) | 2532 ± 24 (2518–2542) | 1678 ± 16 (1669–1685) |
| low pH | T+2 | 07/07/2025 | 7.55 (7.52–7.58) | 30.26 ± 0.77 (30.18–30.8) | 2580 ± 7 (2578–2582) | 2461 ± 7 (2459–2464) | 1671 ± 4 (1669–1673) |
| low pH | T+4 | 07/07/2025 | 7.54 (7.53–7.57) | 32.23 ± 0.35 (31.95–32.47) | 2604 ± 11 (2598–2608) | 2473 ± 10 (2467–2476) | 1694 ± 7 (1690–1696) |

### Biomass and metabolism

The wet weight of each individual was measured at the start of the experiment and every two weeks to quantify biomass change. For each measurement, samples were removed from the aquaria, gently blotted on both sides with paper towels in a standardized manner, and weighed on a precision balance. Photographs were taken at each time point to document algal condition. After weighing, individuals were returned to their respective tanks. The weight of the attached tag was subtracted for the CO_2_ vents individuals.

In addition to biomass change, the effects of temperature and pH on metabolism were assessed through 80-minute incubations. Three replicates per treatment were placed in hermetic 1.3 L transparent containers, each equipped with a small submersible pump to ensure homogeneous water mixing. Each incubation consisted of 40 minutes under artificial light (Mitras Slimline Skywhite) to measure net primary production, followed by 40 minutes in darkness to measure dark respiration. Dissolved oxygen concentrations were recorded using miniDOT Clear Loggers (PME, 0.001 mg/L). Incubations were conducted during the first and final weeks of the experiment to evaluate temporal changes in seaweed metabolism. Furthermore, photosynthesis-irradiance (P-I) curves were performed at the end of the experiment for all treatment combinations, except for vent-origin under present-day pH, which were excluded due to time constraints (n = 3 per treatment). To create the P-I curves, the samples were exposed to 6 sequential irradiances ranging from the minimum to the maximum output of the artificial light (0, 21.8, 47.9, 60.3, 92.5, and 152.1 µmol photons m^-2^ s^-1^), with each irradiance step held for 20 min. The protocol initiated with a 20-minute dark period, followed by the progressive light increments.

### Carbon and nitrogen content

To assess potential long-term shifts in tissue stoichiometry driven by chronic exposure to elevated CO_2_ concentrations, total carbon (C) and nitrogen (N) contents were analysed in individuals remaining at the end of the experimental period (n = 83). Samples were oven-dried at 60 °C and subsequently pulverized into a fine, homogeneous powder using a mortar and pestle. The pulverized samples were then weighed to the nearest 0.001 mg using a Class I precision microbalance and encapsulated in tin cups. Elemental analysis was performed using a FlashEA 112 elemental analyzer (ThermoFinnigan). Samples underwent automated flash combustion at 1020 °C within a quartz reactor tube packed with tungsten trioxide (WO_3_) and copper catalysts.

#### (ii) Herbivory on *Ericaria balearica* across sea urchin’s thermal regime under contrasting pH conditions

To assess how OW and OA may alter herbivory pressure in temperate marine forests, consumption experiments were carried out using a common herbivore of the Mediterranean Sea, *Paracentrotus lividus.* Two complementary sets of laboratory experiments were conducted using independent cohorts from the same target population.

Sea urchins (test diameter 3–4 cm) and *E. balearica* individuals were collected by SCUBA diving from a location (41°57′50″ N, 3°13′42″ E) close to the CEAB-CSIC laboratory. As these experiments focused on the effects of OW and OA on herbivory rather than population-specific differences, locally sourced *E. balearica* individuals were used to avoid additional pressure on the Panarea populations and to maintain a common geographic origin for the urchins and seaweed. Upon arrival, individuals were acclimated to laboratory conditions in temperature-controlled water baths, increasing the temperature at a rate of 0.6 °C Day⁻¹ until treatment temperatures were reached. Low-pH seawater was prepared in header tanks by bubbling CO₂ gas while present-day pH seawater was supplied from separate header tanks. Seawater from each header tank was delivered to the aquaria via individual tubes, with the water flow renewing daily. To minimize variability in hunger state, individuals were not fed for 3 days prior to trials. The first experiment used a factorial design crossing six temperature levels (15, 18, 21, 25, 27, and 29 °C) with two pH treatments set to pH_T_ 7.9 (present-day) and pH_T_ 7.60 (low pH), resulting in 12 treatment combinations. Each treatment consisted of n = 3 aquaria (each aquarium containing a single urchin and a pre-weighed piece of *E. balearica*) and 1 algae-only control aquarium to account for autogenic biomass changes, totaling 48 aquaria (36 treatment + 12 control units). Replication was intentionally conservative to minimize animal collection for conservation and ethical considerations.

The second experiment was designed to refine the thermal performance curve under present-day pH conditions and increase the resolution of the results, as results of the first herbivory experiment clearly showed that OA reduced consumption and flattened the response of *P. lividus.* To maximize statistical power and fine-tune resolution around critical thermal thresholds, the temperature matrix was modified to better capture critical inflection points (15, 18, 21, 23, 26, and 29 °C). The algae-only controls were placed directly inside each experimental aquarium within a small, attached mesh container that allowed water flow but prevented urchin grazing. The experiments were carried out in Spring 2025 and 2026 and lasted 5 and 2 days respectively, depending on the food availability during each trial.

### Field surveys for benthic communities’ characterization

#### (iii) Associated macroalgal community

Benthic community composition and macroalgal biomass under CO₂ vents and ambient pH conditions were assessed during an expedition to Panarea in June 2022. Benthic assemblage samples were collected by scraping the substrate (scrapped area ranging from 1,014 to 1,792 cm^2^). In the laboratory, samples were fully sorted or, for larger samples, a subsample comprising >25% of the total biomass was sorted. Algae were identified to the lowest possible taxonomic level. Biomass was measured as wet weight and as dry weight after oven drying at 65 °C for 48 h.

#### Benthic cover

During the 2025 expedition to Panarea, 3-m long transects were randomly placed at both vent (2 transects) and off-vent (3 transects) sites. Along each transect, benthic cover was assessed every 10 cm using the line-point-intercept method. At each point, the organism or substrate directly beneath the ruler (e.g., identified algal species, dead matte, bare rock, gravel) was visually identified and recorded. When field identification was uncertain, a sample was collected for confirmation under a stereomicroscope in the laboratory. Species were identified to the lowest possible taxonomic level, with the data expressed as the percentage occurrence of each category across all survey points per transect.

#### Data analyses

Biomass changes were analysed by modelling log-transformed mass as a function of time (days since first weighing), temperature (present-day, T+2 °C, T+4 °C), pH (present-day, low), and origin of population (vent, ambient), including all time × factor interactions to test treatment effects on growth rates. Individual identity was included as a random intercept, with a first-order autoregressive [AR(1)] residual correlation structure to account for temporal autocorrelation among repeated measurements of the same individual.

Water bath and aquarium were also tested as random effects to account for potential non-independence arising from shared experimental conditions, but their inclusion did not improve model performance and they were excluded from the final model. Interaction terms were evaluated by likelihood-ratio tests under Maximum Likelihood, and the final model was refitted with Restricted Maximum Likelihood (REML). Growth rates were estimated from the slopes of log-transformed biomass over time using the emtrends () function from R package emmeans and expressed as percent change over14-day intervals. For the Temperature × Population interaction, marginal means were estimated for each population within temperature at representative days, with Tukey-adjusted contrasts.

For productivity measurements, dissolved-oxygen time series were fit for each replicate to obtain rates of change in oxygen concentration (mg O₂ L⁻¹ min⁻¹). These rates were converted to oxygen production or consumption rates based on chamber volume and subsequently normalized by algal biomass to obtain mass-specific net photosynthesis (light periods) and respiration (dark periods). Net photosynthesis was analysed with a mixed model including Period × Temperature × pH × Population and a random intercept for algae ID. Dark-period respiration showed heteroscedasticity, so we used Generalized Least Squares (GLS) with temperature-specific residual variances. Effects were summarized using Estimated Marginal Means (EMMs) and appropriate multiple-comparison adjustments.

Photosynthesis-Irradiance curves were characterized by measuring mass-specific hourly oxygen flux rates (mg O_2_. g^-1^h^-1^) calculated from the linear slopes of dissolved oxygen over time across irradiance intervals (0, 21.8, 47.9, 60.3, 92.5, and 152.1 µmol photons m^-2^ s^-1^). Metabolic parameters were extracted following standard protocols: photosynthetic efficiency (*α*) was calculated separately for each incubation as the slope of a linear regression fitted to the initial linear portion of its P-I curve, identified individually by visual inspection, the light compensation point (I_c_) was determined as the x-intercept of this linear region, maximum photosynthesis (P_max_) was defined as the highest recorded P_net_, and the saturation index I_k_ was calculated as P_max_/α.

Variations in macroalgal tissue nutrient content (%C, %N, and C:N) were analyzed using Generalized Linear Models (GLMs) with a Gaussian distribution and an identity link function. Models included population, pH, temperature, and all two– and three-way interactions as fixed factors. Statistical significance was evaluated using Type-III Analysis of Variance (ANOVA). Residual normality and homoscedasticity assumptions were verified via diagnostic plots.

The effects of OA x OW on sea urchin feeding pressure were evaluated by analyzing consumption rates (g WW day⁻¹) using a GLM with a Gaussian error distribution and an identity link function. Temperature was incorporated as a second-order polynomial to capture the non-linear geometry of the thermal response. Global significance testing was performed using a Type III ANOVA with Wald F-tests to account for the differences between the 2025 and 2026 datasets. Finally, specific effects of OA across the thermal gradient were assessed using post-hoc pairwise contrasts based on EMMs.

To assess differences in the associated macroalgal communities, biomass was standardized by substrate area (g cm⁻²). Two subsampled records were back-calculated to whole-sample biomass. Community structure was visualized with NMDS based on Bray–Curtis dissimilarities of log1p-transformed biomass density and tested with PERMANOVA (999 permutations; model: community ∼ station). Homogeneity of dispersion was assessed with PERMDISP, and SIMPER identified taxa contributing most to between-station differences.

To measure benthic cover, point counts were summed by species group and converted to percent cover. Treatment-level differences were summarized by mean and standard error. Differences in overall benthic community composition between pH treatments were evaluated with PERMANOVA (Bray–Curtis; 999 permutations) and checked for dispersion differences with PERMDISP.

All analyses were conducted in R (v4.5.3, R Core Team, 2026; www.r-project.org) using the packages *nlme* (Pinheiro et al., 2025) and *glmmTMB* (Brooks et al., 2017) for mixed-effects modelling, *emmeans* (Lenth & Piaskowski, 2026) for estimated marginal means and post-hoc contrasts, *car* (Fox & Weisberg, 2019) for Type III ANOVA, *vegan* (Oksanen et al., 2026) for multivariate community analyses, and *seacarb* v3.3.4 (Gattuso et al., 2026) for carbonate-system calculations. Data processing and visualization were performed using *tidyverse* (Wickham et al., 2019).

## Results

### Seaweed performance

Seaweed biomass declined over time in all treatments, and only temperature altered the rate of loss (Fig. 2). Mixed-effects slopes showed a faster decline at T+2 °C than at T+4 °C (difference = −0.0105 log-biomass day⁻¹ Tukey-adjusted p = 0.0011), whereas the present-day temperature slope did not differ from either warm level (p = 0.12 and 0.24). Consistent with these trends, mean biomass did not differ among temperatures at the start, but at the endpoint T+2 °C < T+4 °C (Tukey p = 0.013); the present-day treatment did not differ from either warm level (p ≈ 0.31–0.32). No pH effect was detected on biomass change. The interaction between population and temperature had a marginal effect on biomass change (p = 0.0781). Nonetheless, post-hoc contrasts at present-day temperature indicated that vent samples retained more biomass than ambient-site samples by mid-to late experiment (day 15 p = 0.035; day 28 p = 0.0077; day 42 p = 0.0024; day 48 p = 0.0019). At day 48, the vent population had roughly double the biomass of the ambient population. By contrast, at T+2 °C and T+ 4 °C the two populations did not differ at any checked day (all p > 0.5).

**Figure 1.**
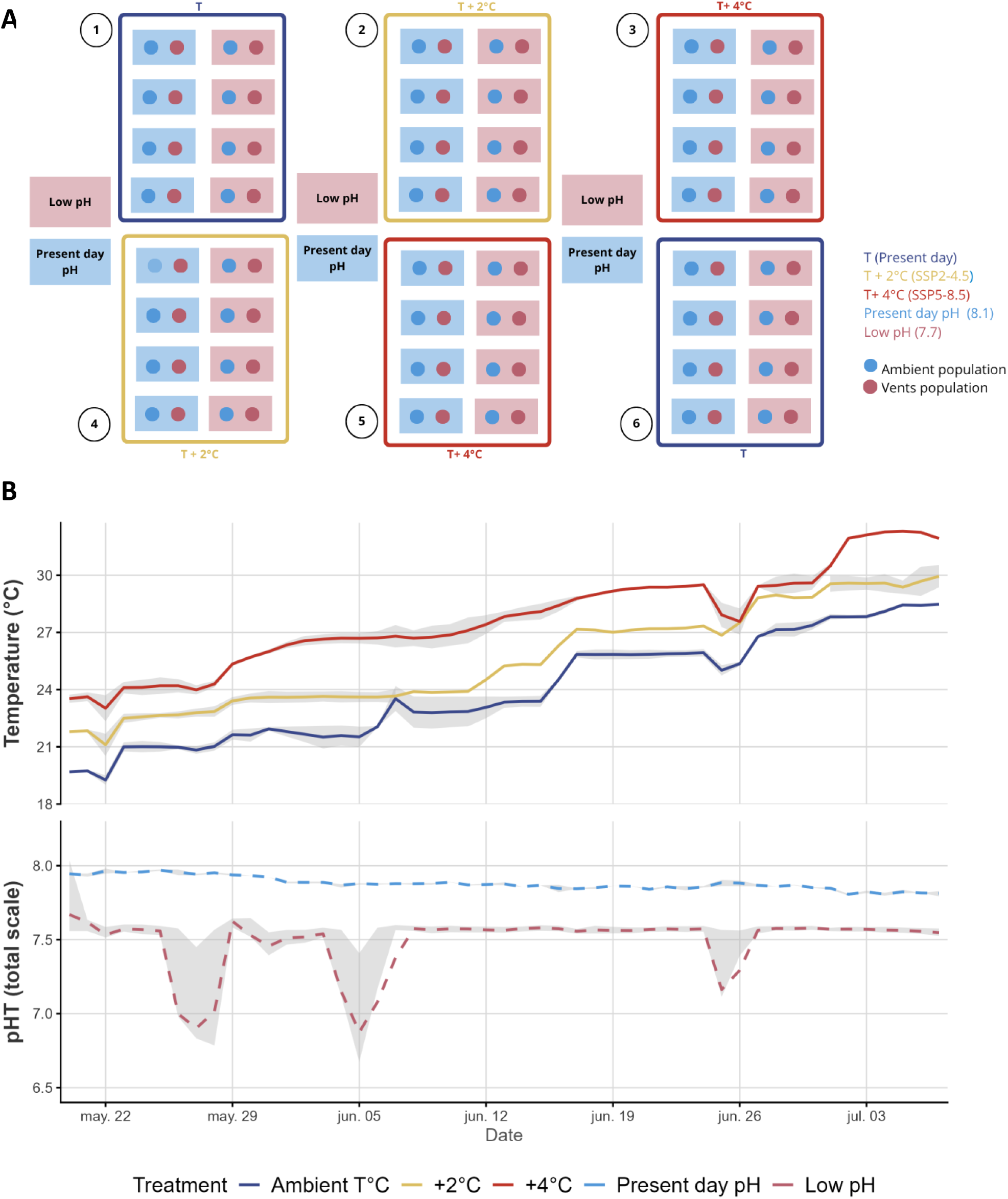
Experimental set-up for the physiological experiment on *E. balearica*. (A). Six water baths held individual aquaria allocated to crossed pH_T_ (present-day, low) and temperature (present-day, T+2 °C, T+4 °C) treatments. Colors indicate treatment assignments. Daily temperature and pH_T_ trajectories across experimental treatments (B). Solid lines represent temperature trajectories, while dashed lines indicate pH treatments. Shaded areas indicate ± SE for temperature and the 25th–75th percentile range for pH_T_. Values were recorded every minute using temperature and pH sensors connected to a GHL Profilux controller. Two brief power outages during the experiment caused short interruptions in temperature control, and CO₂ delivery failed on three occasions, temporarily lowering pH in the low-pH treatments.

**Figure 2.**
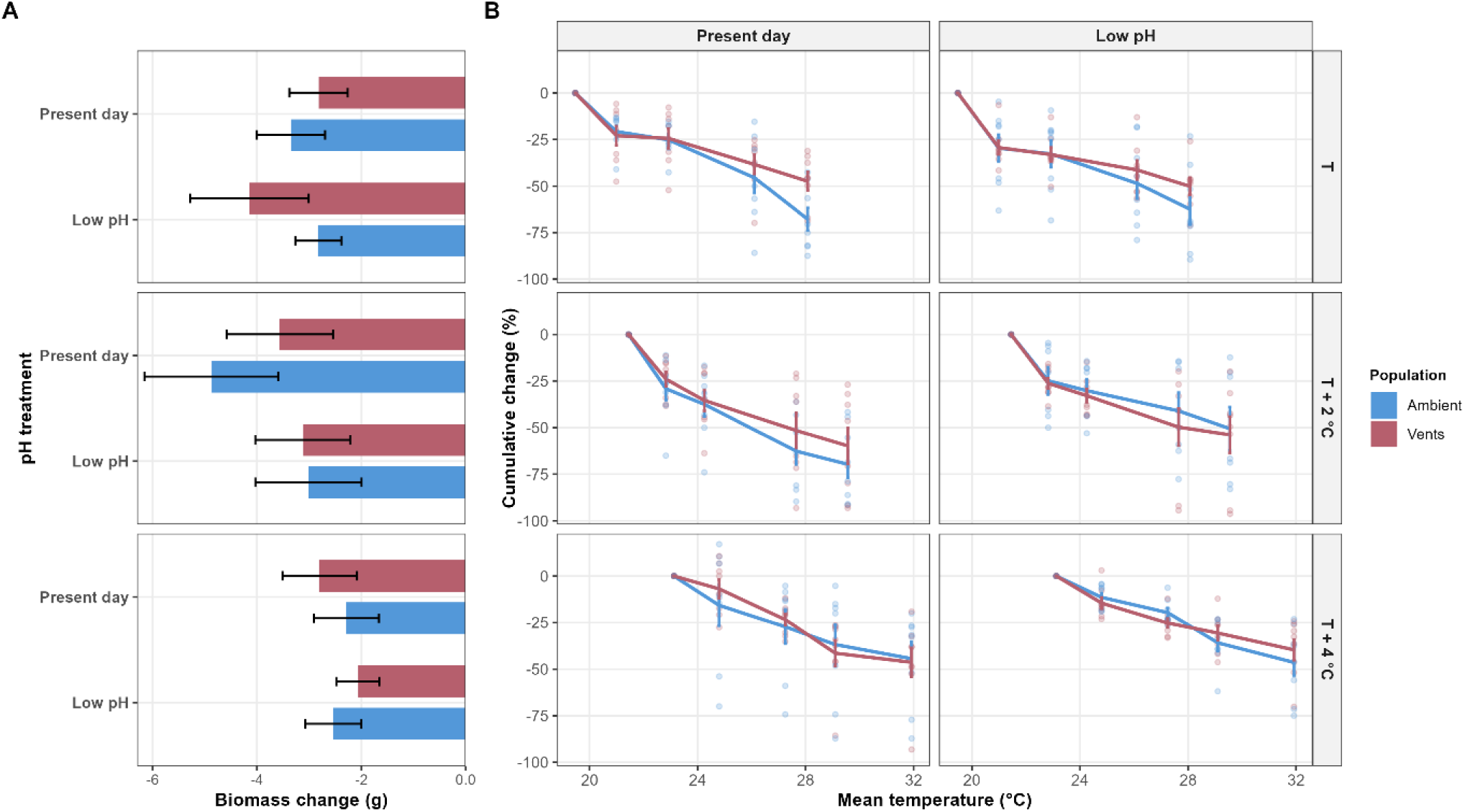
(A) Mean change in biomass (g) between initial and final measurements for each temperature × pH treatment and population. (B) Mean cumulative biomass change (%) over the experimental period, plotted against the mean temperature recorded during each measurement interval (14 days). Points represent individual replicates, while lines and error bars show treatment means ± SE.

Linear mixed-effects modelling revealed a significant interaction between incubation period, temperature, and pH on macroalgal net photosynthesis (p < 0.001). Population origin showed no significant main or interactive effects (p > 0.12), and post-hoc comparisons were therefore averaged across populations. Initial NP rates did not differ significantly among treatments (Tukey p > 0.53). NP declined overall (Period, p < 0.0001), although the magnitude varied with temperature and pH. The largest estimated decline occurred under low pH at present-day temperature (Holm p = 0.0027). At the final incubation, NP was significantly higher under T+2 °C and T+4 °C than present-day temperature only under low pH (Tukey p < 0.0001). Low pH reduced final NP at present-day temperature (Holm p < 0.0001), but increased it at T+2 °C (p = 0.0116). Dark respiration also exhibited a significant Period × Temperature × pH interaction (p < 0.0001), with respiratory oxygen consumption increasing significantly over time under low pH at present-day temperature (Holm p < 0.0001; Fig. 3). Gross primary production (GPP), showed contrasting trends under low pH, increasing between initial and final incubations at T+2 °C but declining at T+4 °C in both populations. Three final incubations yielded negative estimates, indicating greater oxygen consumption under illumination than in darkness rather than negative gross photosynthesis. These patterns suggest that reduced production may contribute to NP declines in some treatments, although interpretation is limited by low replication.

**Figure 3.**
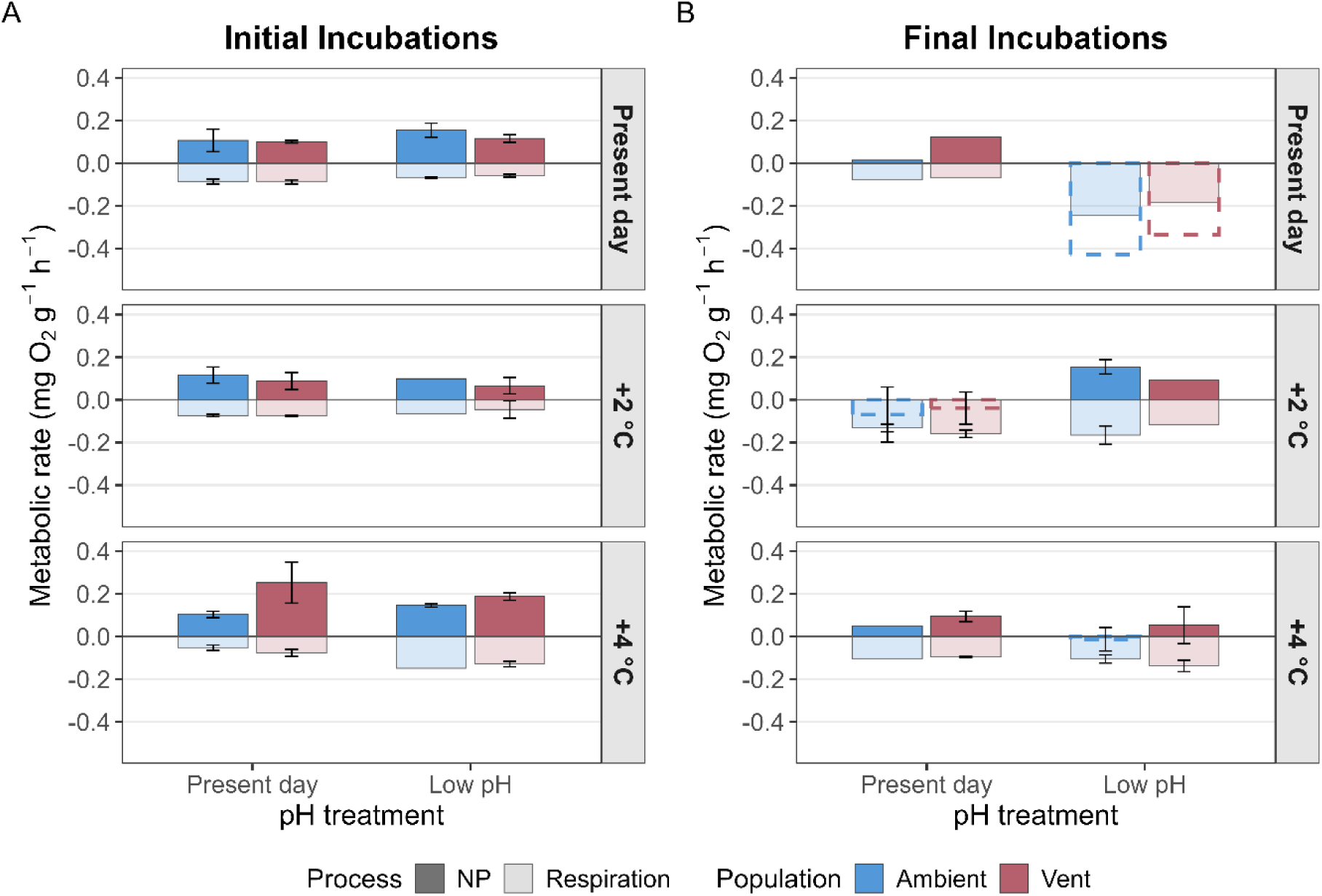
Mean metabolic rates of ambient and vent populations across temperature (present-day, T+2 °C, T+4 °C) and pH treatments (Present day pH_T_ 7.9 and low pH_T_ 7.6). (A), Initial incubations conducted on 21-22 May. (B), Final incubations conducted on 25-27 June. Net photosynthesis (NP) is shown in darker shades, with negative values represented by dashed outlines, whereas dark respiration is shown in lighter shades. Bars represent mean metabolic rates, and error bars indicate ± SE.

For the P-I curve analysis, neither temperature nor pH significantly affected photosynthetic efficiency (α), light compensation point (Ic), maximum photosynthesis (Pmax) or saturation irradiance (Ik) in ambient-origin individuals (n = 8; all p > 0.05). Mean α under present-day pH was 0.00278, 0.00184 and 0.00176 mg O₂ g⁻¹ h⁻¹ per µmol photons m⁻² s⁻¹ at present-day, T+2 °C and T+4 °C, respectively, compared with 0.00268 and 0.00218 mg O₂ g⁻¹ h⁻¹ per µmol photons m⁻² s⁻¹ under low pH at T+2 °C and T+4 °C. Under low pH (*n* = 6), no significant effects of population origin or temperature were detected for any parameter (all *p* > 0.05). Descriptively, α was higher in vent-origin than ambient-origin individuals at T+2 °C (0.00383 vs. 0.00268 mg O₂ g⁻¹ h⁻¹ per µmol photons m⁻² s⁻¹) and T+4 °C (0.00268 vs. 0.00218 mg O₂ g⁻¹ h⁻¹ per µmol photons m⁻² s⁻¹), while *Pmax* declined with warming in ambient-origin individuals under present-day pH (0.211, 0.117 and 0.048 mg O₂ g⁻¹ h⁻¹ at present-day, T+2 °C and T+4 °C, respectively); however, neither pattern was statistically significant.

Nutrient analysis revealed no significant interactive effects of population, pH, and temperature on the C:N ratio of the macroalgae (p > 0.05 for all interaction terms) nor any significant effect of pH (p = 0.545). However, marginal effects were observed for temperature (p = 0.055) and population (p = 0.097). Biologically, the tissue C:N ratio increased progressively with rising temperatures, reflecting the depletion of relative nitrogen content and a subsequent decline in macroalgal nutritional quality under thermal stress.

### Herbivory on Ericaria balearica across sea urchin’s thermal regime under contrasting pH conditions

Sea urchin consumption rates were significantly affected by seawater pH (p = 0.034), with a marginally significant interaction between pH and temperature (p = 0.054). Under present-day pH, a classical thermal performance curve was observed, with consumption peaking around 22 °C and decreasing toward thermal limits (Fig. 4). In contrast, the GLM showed a clear trend toward the flattening of this curve under ocean acidification conditions. Pairwise post-hoc contrasts confirmed that the impact of low pH was temperature-dependent; feeding rates did not differ at the cold extreme of 15 °C (p = 0.615), but low pH caused a severe reduction in consumption across the entire intermediate and optimal thermal window from 18 °C to 27 °C (p < 0.001 for 18–26 °C; p = 0.002 at 27 °C). The maximum suppression of herbivory occurred at the operational thermal optimum (21–23 °C), where low pH reduced estimated feeding rates by 90.6% and 90.3%, respectively, relative to present-day conditions. Finally, at the upper thermal limit of 29 °C, consumption was low in both treatments, eliminating any remaining differences between pH levels (p = 0.642).

**Figure 4.**
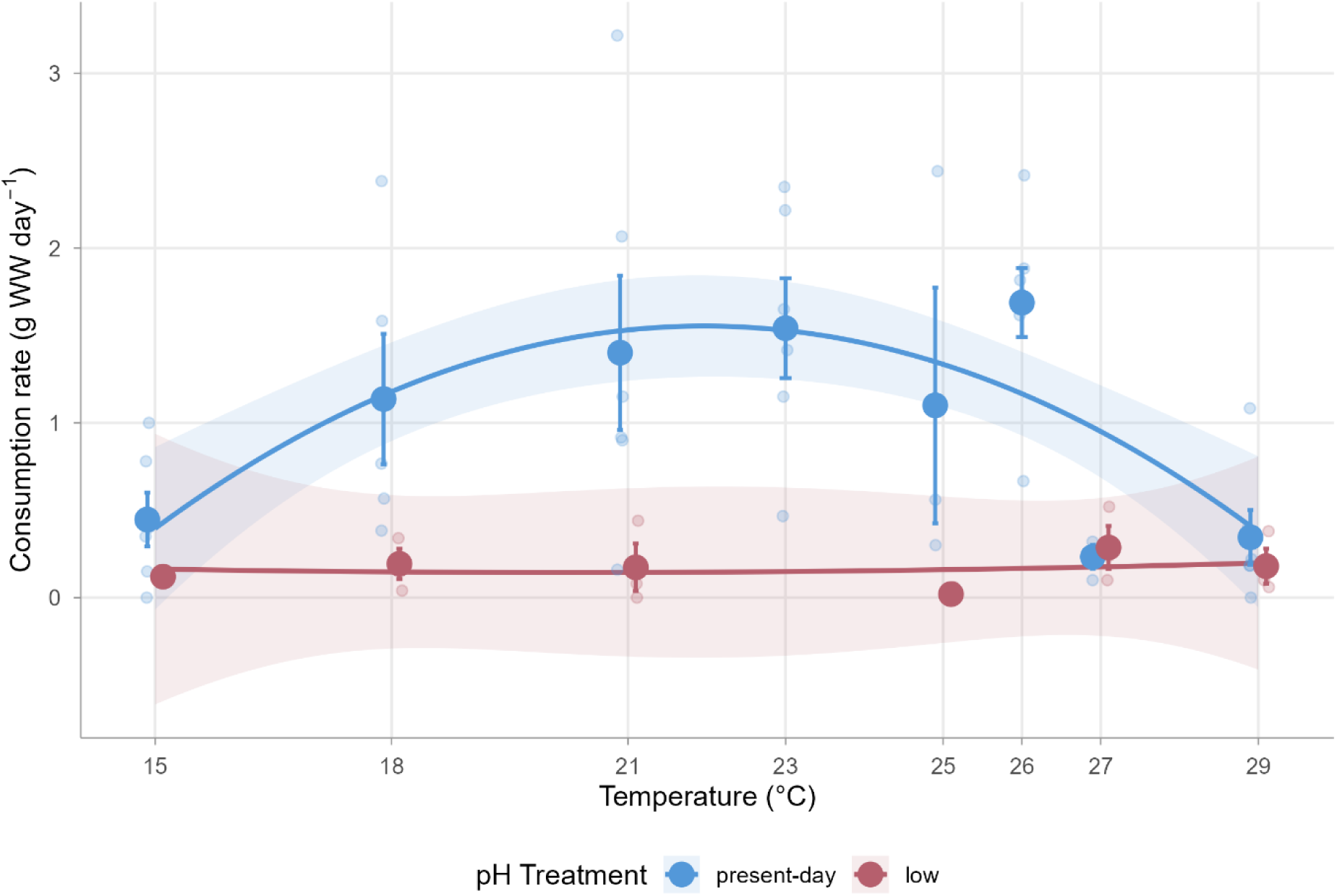
Consumption rates (g WW day⁻¹) of *P. lividus* feeding on *E. balearica* across 8 temperature treatments and 2 pH levels (present day pH_T_ 7.9, low pH_T_ 7.6).

### Associated macroalgal community

The community composition of *E. balearica* forest scrapings differed strongly between ambient– and low-pH sites (PERMANOVA, p = 0.005). SIMPER analysis indicated that *E. balearica* was the main contributor to the observed dissimilarity, accounting for 65.0% of the total dissimilarity, followed by filamentous Rhodomelaceae (14.2%), and *Codium bursa* (7.2%) which were more abundant under low pH (fig.5).

**Figure 5.**
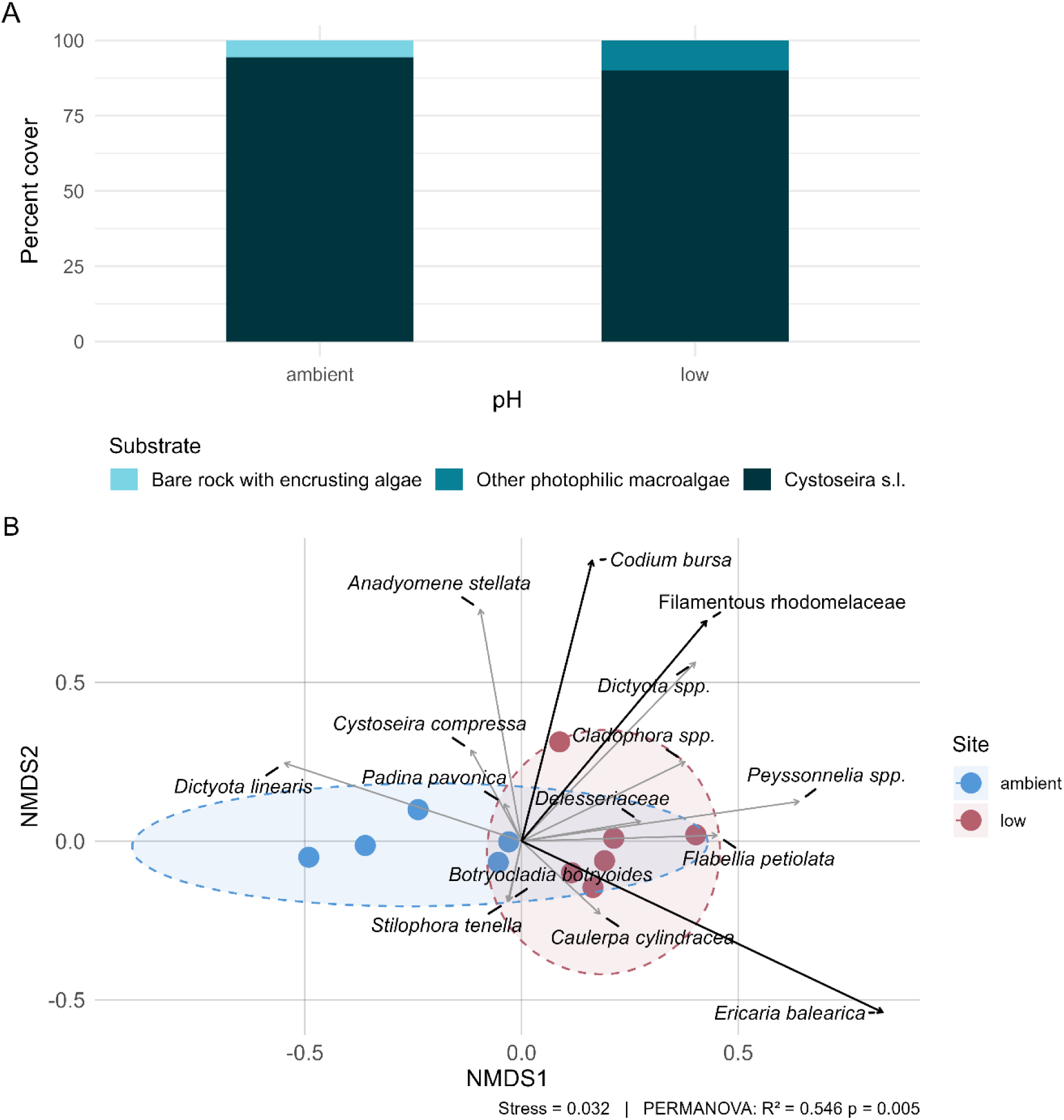
Species frequency and biomass at ambient– and low-pH sites (Panarea, Italy). (A) Species frequency based on point-contact transects. (B) NMDS of species biomass from *E. balearica* forest scrapings; Arrows indicate directions of increasing species biomass, with their lengths reflecting the strength of association with the ordination. Black arrows indicate significant associations (*p* < 0.05), while grey arrows indicate non-significant associations.

The point contact data show that *Cystoseira s.l.* cover was high in both present-day and low pH sites (94% and 90%, respectively). A quasibinomial GLM detected no significant effect of pH on *Cystoseira s.l.* cover (p = 0.20), indicating that this canopy-forming macroalga remains dominant under reduced pH conditions at the scale of our transects.

## Discussion

Our findings suggest that the future of marine forests under climate change will be primarily driven by shifts in community-level processes rather than at the population level linked to changes in physiology responses. Although photophilic canopy-forming macroalgae seem to respond positively to progressive OA—benefiting from chronic exposure to elevated *p*CO₂—this advantage is minor compared to the expected changes in herbivory pressure and shifts in species composition. OW directly affected *E. balearica* performance by accelerating the senescence period, while the interaction between OA and OW altered herbivory pressure. Any beneficial effects of enhanced performance under OA appear to accumulate at the community scale, yielding a slight biomass increase in *E. balearica* locally acclimated to low pH, alongside simultaneous reorganizations of the understory community driven by species-specific pH tolerances. Ultimately, these concurrent community-wide disruptions seem to dictate the long-term trajectory of the ecosystem.

### Growth: temperature, pH and phenology effects

The experimental results indicate that the individuals from CO_2_ vents marginally benefited from reduced biomass loss at present-day temperatures, though this advantage was masked under elevated temperature regimes. Overall, pH had no significant main effect on biomass trajectories, suggesting that elevated *p*CO₂ does not exert an acute impact; rather, performance differences are likely tied to the long-term benefits of chronic exposure or local adaptation. This origin-specific pattern disappeared under warming (T+2°C and T+4°C), which accelerated summer conditions and senescence across all treatments. Consequently, while population origin can modulate performance under contemporary temperatures, warming rapidly erodes this buffering capacity. Finally, OW and OA exhibited no interactive effects—whether additive, synergistic, or antagonistic—on biomass dynamics.

The general biomass loss can be explained by the timing of the experiment and the species phenology. While these types of experiments often result in stressful conditions for seaweed compared to natural conditions, fucalean canopies are strongly seasonal in the Mediterranean. For *E. balearica* specifically, observations at Calvi (Corsica) documented two growth periods, with most biomass produced during winter–spring and a shorter autumn phase, branch shedding occurred in both late summer and winter (Hoffmann et al., 1992). Regional syntheses for the western Mediterranean (Balearic Islands, SE Spain) similarly report winter–spring growth, a summer resting phase, and autumn renewal, with some geographic shifts in timing (Gomez Garreta et al., 1982; Luque & González, 2004). In Mallorca, by early summer *E. balearica* loses most primary and secondary branches and thalli are reduced to basal parts; new axes appear in October and by November plants resemble the winter form (Gómez et al., 1982). Our experiment therefore overlapped the late-summer senescence/resting phase, which likely dampened treatment contrasts with T+2°C and T+4°C reaching these temperatures earlier in the experiment than present-day temperature treatments and being exposed to senescence temperatures for longer periods during the experiment. During senescence, reduced tissue production and the loss of photosynthetic branches may limit the extent to which enhanced inorganic carbon availability under OA is translated into biomass accumulation, potentially masking OA effects on growth and metabolism. Conversely, during the active growth period, when photosynthetic activity and tissue production are higher, increased CO₂ availability may have a greater potential to stimulate carbon assimilation and biomass production (Ní Longphuirt et al., 2013; Baggini et al., 2014; Graiff et al., 2015). Repeating the experiment across contrasting phenological periods would therefore help determine whether there are any interaction effects of OA and OW on *E. balearica* performance and if they are seasonally dependent. On that note, we recognize that sampling *E. balearica* before the growth period (in winter), especially in remote places such as Panarea, is challenging and many studies concentrate in this window for the same reasons.

### Short-term metabolism

Short-term metabolic responses were consistent with the observed biomass trajectories, although the underlying processes varied among treatments. At present-day temperature and low pH, declining NP coincided with increased dark respiration, whereas under T+4°C and low pH, reduced GPP without a consistent increase in respiration suggests that lower photosynthetic activity may also have contributed. As experimental temperatures progressively increased towards summer conditions, senescence and the loss of photosynthetic branches may have constrained oxygen production, potentially masking direct effects of warming and elevated CO₂ on metabolism.

Although the P–I curve analyses detected no significant treatment effects, vent-origin individuals showed a tendency towards higher photosynthetic efficiency under low pH at both warming levels, together with higher *Pmax* at T+2°C. These trends should be interpreted cautiously given the considerable inter-individual variability and limited replication in some treatments. Photosynthetic performance, particularly *Pmax*, can vary considerably among Mediterranean Fucales depending on species, depth and season (Sant & Ballesteros, 2021). The maximum irradiance achievable with our lamps (152 µmol photons m⁻² s⁻¹) fell within the approximate saturation irradiance (*Ik*) range reported for Fucales (120–200 µmol photons m⁻² s⁻¹; Ballesteros & Sant, 2022). Thus, this irradiance may have been sufficient to approach saturation in some individuals, although additional measurements at higher irradiances would be needed to confirm the plateau across all P–I curves.

### Herbivory: acidification minimizes top-down control

Our findings show that herbivory is significantly reduced under low-pH conditions, a response that is highly consistent with the known sensitivity of echinoderms to acidification. Because echinoderms possess high-magnesium calcite skeletons, they are frequently identified as key taxa vulnerable to OA (Dupont et al., 2010). This vulnerability is heavily mirrored by field observations at natural volcanic CO_2_ vents, where sea urchin abundances drop sharply as pH declines (Hall-Spencer et al., 2008; Johnson et al., 2012; Coni et al., 2021; Teixidó et al., 2018). Mechanistically, acidification can impair calcification, disrupt acid–base regulation, and hinder feeding performance, culminating in a clear relaxation of top-down grazing pressure (Hall-Spencer et al., 2008; Teixidó et al., 2018).

In Mediterranean rocky reefs, this reduction in echinoderm grazing has profound ecological implications. Sea urchins, primarily *Paracentrotus lividus,* represent the dominant herbivore pressure in these ecosystems, capable of driving regime shifts from complex macroalgal forests to low-productivity, simplified alternative stable states known as urchin barrens (Sala et al., 1998; Filbee-Dexter & Scheibling, 2014; Boada et al., 2017). While temperature typically shapes herbivory—with *P. lividus* grazing rates known to peak around 21°C (Pagès et al., 2018)—our results reveal that low pH alters this thermal relationship. Specifically, acidification not only reduced overall *P. lividus* consumption but also largely flattened its thermal response across the tested range, erasing any identifiable optimal temperature for herbivory. Under OW, grazing may likewise decline once temperatures approach or exceed the species’ upper thermal limits, as thermal stress increasingly constrains urchin performance (Pagès et al., 2018; Jiménez-Ruano et al., 2026). Recent work on barren-forming sea urchins further indicates that warming can constrain urchin performance near the warm edge of their thermal distributions, although the magnitude and direction of responses vary among species and populations (Butler et al., 2026a, b, c). Thus, progressive warming may initially maintain or increase grazing within the species’ thermal performance range, but could ultimately weaken *P. lividus* herbivory as temperatures move beyond its optimal range.

While barren states are typically reinforced by high urchin recruitment and a lack of predators, our results suggest that chronic OA, together with temperatures approaching the upper thermal limits of *P. lividus*, could weaken the grazing pressure required to initiate or maintain overgrazing. In this context, both stressors may relax urchin-mediated top-down control, although through different pathways: OA directly reduced consumption across the thermal gradient in our experiment, whereas OW is expected to produce a non-linear response in which grazing declines only as temperatures exceed the species’ thermal optimum. The combined effects of these stressors could therefore alter the strength and timing of herbivore control on Mediterranean macroalgal forests.

However, the wider herbivore group displays highly variable responses to shifting climate drivers, indicating that reduced grazing by *P. lividus* will not necessarily translate into a proportional decline in overall top-down control. For instance, the herbivorous fish *Sarpa salpa*—which heavily influences the biomass and vertical distribution of *Cystoseira s.l.* species (Vergés et al., 2009; Gianni et al., 2017)—appears to show increased overall herbivory impact on *Posidonia oceanica* at CO_2_ vents with low-pH conditions (Mirasole et al., 2026).

OW may further alter the composition of the herbivore assemblages by accelerating the tropicalization of the Mediterranean and facilitating the range expansion and recruitment of thermophilic and tropical vagrant fishes (Vergés et al., 2014). As temperatures rise, these fish consumers can become key drivers of macroalgal consumption, potentially shifting the ecosystem from an urchin-dominated grazing regime to a fish-dominated one. Thus, although OW and OA may reduce grazing by *P. lividus* when environmental conditions exceed its optimal range or become physiologically stressful, other herbivores may increase in abundance or maintain their feeding activity under the same conditions.

Consequently, future changes in grazing pressure will depend not only on how climate change affects the feeding rates of individual consumers, but also on which herbivore populations persist and become abundant under altered environmental conditions. The expected reduction in *P. lividus* grazing under acidification and warming, at sufficiently high temperatures, could therefore be partly offset by shifts toward other herbivores, particularly fishes favored by Mediterranean warming. The resulting balance between herbivory and macroalgal persistence will ultimately depend on how these changing consumer assemblages track environmental conditions, with implications for whether canopy-forming macroalgal forests are maintained or replaced by more simplified, turf-dominated states.

### Community composition

*E. balearica* canopy cover was very similar between ambient pH and CO_2_ vents conditions. While marine forests showed signs of a healthy state at both sites, we identified a slight increase in biomass under low pH. This suggests that the positive trends observed in our laboratory experiments can scale up to the ecosystem level, with OA potentially enhancing the performance of populations chronically exposed to elevated *p*CO_2_. A plausible mechanism driving this pattern is physiological acclimatization, which involves plastic shifts in carbon uptake and photophysiology under elevated CO₂, and has been reported for some fucoids and other brown algae (Celis-Plá et al., 2017; Hurd et al., 2020). However, our experimental results indicate that this advantage is context-dependent and heavily constrained by temperature, manifesting only under ambient thermal regimes. Because factors like light and nutrient availability further modulate the direction and magnitude of carbon fertilization, any vent-related benefit is unlikely to be universal (Koch et al., 2013; Hurd et al., 2020). Overall, the observed changes suggest a reorganization of the forest-associated assemblage despite similar *E. balearica* canopy between pH conditions. The understory restructuring described is highly consistent with observed global trends. Several of the taxa contributing to the dissimilarity were fleshy, filamentous, or non-calcified, consistent with patterns reported from other Mediterranean CO₂ vents, where low-pH assemblages tend to be characterized by seasonal, fast-growing and non-calcified functional traits (Teixidó et al., 2018; Carlot et al., 2026).

Filamentous Rhodomelaceae, particularly *Vertebrata fruticulosa*, were more abundant under low pH, potentially benefiting from enhanced carbon availability due to their fast-growing, non-calcified morphology which may favour rapid carbon acquisition (Teixidó et al., 2024). Similarly, *Dictyota* spp. increased under low pH, consistent with reports of CO₂-enhanced growth and abundance in some species (Cornwall et al., 2017; Burnham et al., 2022), although *D. linearis* showed the opposite pattern, highlighting species-specific responses. *Codium bursa* also increased under low pH, potentially reflecting indirect effects of acidification, as its high-affinity carbon-concentrating mechanism makes a direct CO₂ benefit less likely (Cornwall et al., 2017). Finally, *Peyssonnelia* spp. were more abundant under low pH, consistent with increased performance under elevated CO₂ reported for the lightly calcified *P. squamaria* (Yıldız, 2018). These patterns suggest that acidification may restructure understory composition through species-specific physiological responses and indirect ecological effects.

When evaluating the overarching future of these ecosystems, the localized beneficial effects of elevated *p*CO_2_ on canopy biomass appear insufficient to build up broad-scale resilience against multifaceted regional disturbances. Local and regional drivers—such as marine heatwaves, habitat degradation, overgrazing, eutrophication, and chronic pollution—remain far more decisive in controlling Mediterranean forest status than gradual, OA and OW alone (Mangialajo et al., 2008; Sales et al., 2011; Boada et al., 2017). Consequently, future projections of marine forest trajectories must transition toward multi-stressor frameworks that integrate seasonally varying thermal constraints, pH-sensitive grazer dynamics, and local habitat quality.

## Acknowledgements

This work was done under the framework of MITnPOINT project funded by the MCIN/AEI/10.130339/5011000011033, FEDER, and 4OCEANS-MOPGA ANR-17-MOPGA-0001 research projects. JB acknowledges the support received by the I+D+i project RyC 2021-033650-I and the European Commission-European Union’s Horizon 2020 MSCA-SHIFT2SOLVE with which the conceptual basis of this work was conceived. JC was supported by a Beatriu de Pinós postdoctoral fellowship (2024 BP 00106) funded by the Government of Catalonia and co-funded by the European Union under the Marie Skłodowska-Curie COFUND programme. Authors are grateful to Lena Saiz, Francesco Martini, Uxue Moreda, Roger Franch, and Andrea Fogliuzzi (Amphibia Diving), the Panarea ECCSEL NatLab-Italy laboratory of OGS and its scientific staff for their support in the laboratory, analyses and field respectively. FORESCUE, (PANAREA 2022 Expedition).

